# Contribution of transition and stabilization processes to speciation is a function of the ancestral trait state and selective environment in *Hakea*

**DOI:** 10.1101/207373

**Authors:** Byron B. Lamont, Sh-hoob M. El-ahmir, Sim Lin Lim, Philip K. Groom, Tianhua He

## Abstract

Currently the origin and trajectories of novel traits are emphasised in evolutionary studies, the role of stabilization is neglected, and interpretations are often *post hoc* rather than as hypothesised responses to stated agents of selection. Here we evaluated the impact of changing environmental conditions on trait evolution and stabilization and their relative contribution to diversification in a prominent Australian genus, *Hakea* (Proteaceae). We assembled a time-based phylogeny for *Hakea*, reconstructed its ancestral traits for six attributes and determined their evolutionary trajectories in response to the advent or increasing presence of fire, seasonality, aridity, nectar-feeding birds and (in)vertebrate herbivores/granivores. The ancestral *Hakea* arose 18 million years ago (Ma) and was broad-leaved, non-spinescent, insect-pollinated, had medium-sized, serotinous fruits and resprouted after fire. Of the 190 diversification events that yielded the 82 extant species analysed, 8–50% involved evolution, stabilization or re-evolution (reversal) of individual novel traits. Needle leaves appeared 14 Ma and increased through the Neogene/Quaternary coinciding with intensifying seasonality and aridity. Spinescence arose 12 Ma consistent with the advent of vertebrate herbivores. Bird-pollination appeared 14 Ma in response to advent of the Meliphagidae in the early Miocene. Small and large woody fruits evolved from 12 Ma as alternative defenses against granivory. Fire-caused death evolved 14 Ma, accounting for 50% of subsequent events, as fire became less stochastic. Loss of serotiny began in the late Miocene as non-fireprone habitats became available but only contributed 8% of events. Innovation and subsequent stabilization of functional traits promoted the overall species diversification rate in *Hakea* by 15 times such that only three species now retain the ancestral phenotype. Our approach holds great promise for understanding the processes responsible for speciation of organisms when the ancestral condition can be identified and the likely selective agents are understood.

## INTRODUCTION

Studies that capture patterns of speciation associated with changes in environmental conditions provide compelling support for the key role of functional trait shifts in the process of evolution by natural selection (Jetz *et al.*, 2012). Natural selection can induce the evolution of novel traits whose fitness exceeds that of the incumbent trait (directional selection) or perpetuation of the current trait whose fitness exceeds that of a former or alternative trait (stabilizing selection) (Lemey *et al.*, 2009). Phylogenetic methods have been developed to investigate a wide range of questions regarding species evolution, including the inference of ancestral traits (He *et al.*, 2011, 2012; Crisp *et al.*, 2011) and to address the relationship between traits and rates of speciation (Litsios *et al.*, 2014). While currently the origin and evolutionary trajectories of novel traits are emphasised the role of stabilization has been neglected and interpretations have often been *post hoc* rather than as hypothesised responses to stated agents of natural selection. This is partly because of ignorance of the advent or strength of the postulated selective agents. Significant questions remain: To what extent do directional and stabilizing process contribute to trait proliferation? Do their contributions vary between attributes, traits and/or over geological time? Is the proliferation of a trait at the expense of its alternative traits? Can patterns of directional and stabilizing selection over time be interpreted in terms of the advent or changes in the intensity of particular agents of selection?

### Theory and concepts

Each alternative state of a species attribute is here termed a trait. Increase in occurrence of a given trait through a phylogeny is defined as trait proliferation (He *et al.*, 2011, Lamont *et al.*, 2013). The fraction of total diversification events that result in the presence of that trait is the trait proliferation rate (this may also be given on an absolute basis per unit time, as for species diversification). Trait proliferation results from two evolutionary processes: transition – a new trait arises during the event, and stabilization – the trait is conserved during the event. Transition rate (TR) is the fraction of events in which the trait arises relative to the maximum number in which that trait could occur, while stabilization rate (SR) is the fraction of events in which the trait is retained relative to the maximum number in which that trait could occur. Thus, the (net) trait proliferation rate (PR) = TR + SR. Generally, for a pair of opposing traits, 1 and 2, evolving in a clade with a total of *n*/2 nodes and thus *n* diversification events for the period of interest, PR_1_ = TR_2/1_ + SR_1/1_, TR_2/1_ = Σ(2**→**1 events)/*n*, and SR_1/1_ = Σ(1**→**1 events)/*n*. Trait reversals are successive transitions that return the phenotype to the previous trait state in the lineage, as an inverse function of the stability through time of the selective pressure for that trait. The concepts of rates of diversification, proliferation, stabilization and transition, and reversals are illustrated with a concrete example in Fig. 1.

**Fig. 1.**
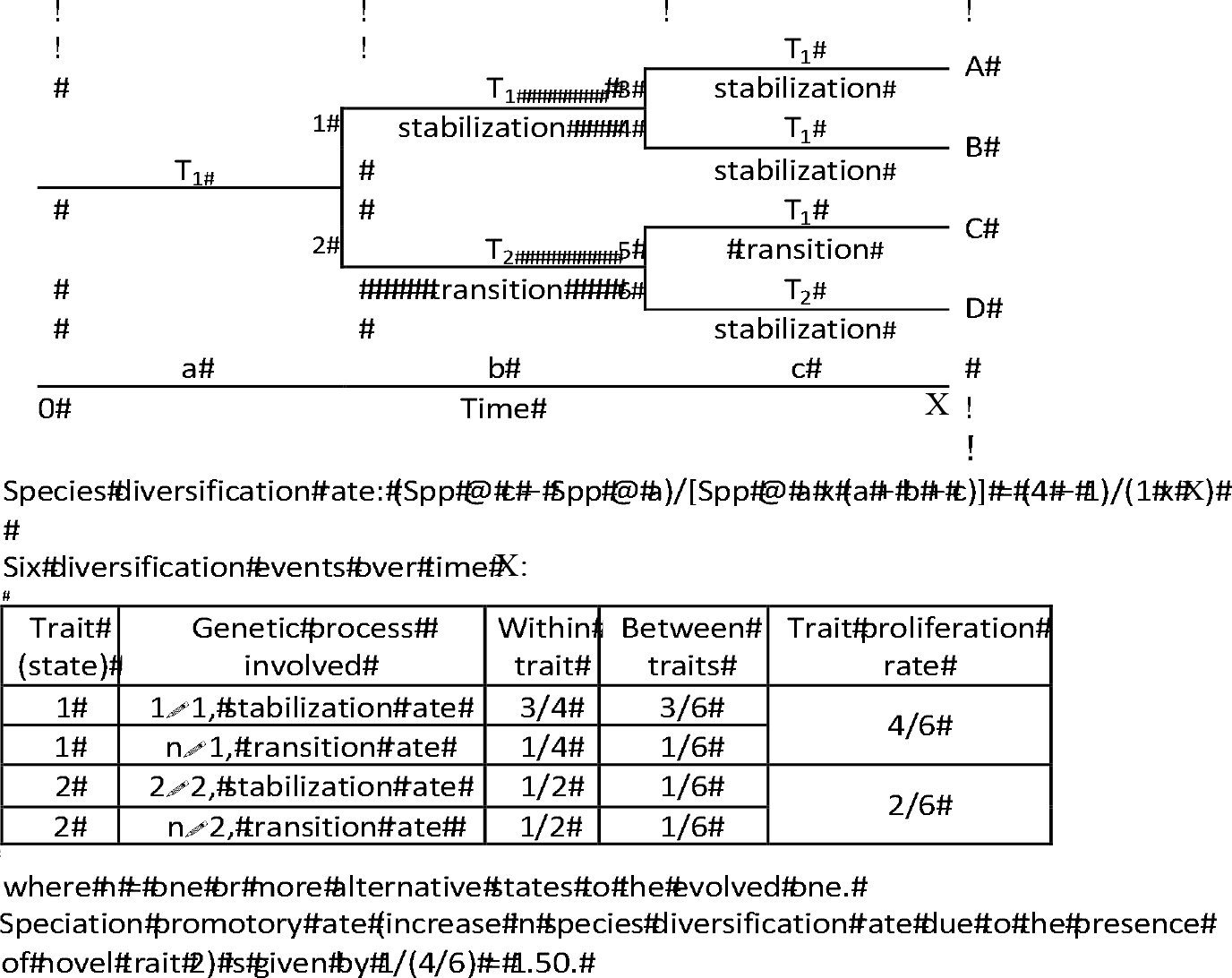
Hypothetical phylogeny showing the evolution of four species from six diversification events with proliferation of two alternative traits, 1 (ancestral) and 2 (novel), of a given character over time, due to both stabilization and transition processes. Diversification rate is relative to the starting number of species/lineages and the time interval, while proliferation, stabilization and transition rates are relative to the maximum number of events in which that trait could occur. Note that the two transitions that yield Sp C is an example of a reversal (to T1).

A new agent of directional selection usually operates in a different spatial or temporal dimension than the existing agents and becomes a supplementary force that initially retards speciation and then promotes it once an adapted genotype has evolved followed by rampant speciation into the “vacant niche” now available as the new trait proliferates (Lamont *et al.*, 2013). The premise here is that the more habitats (niches) a given area can be divided into, the greater the opportunities for novel genotypes to arise. Accepting that new, alternative traits supplement rather than replace ancestral traits as options, the contribution of novel traits to speciation can be calculated as the inverse of the percentage contribution of the ancestral trait to all subsequent diversification events (Y): speciation promotional rate (SPR) = 100/Y. Thus, the smaller Y, the greater the contribution of novel traits to the subsequent diversification events.

### *Evolution and adaptations of* Hakea

Nutrient-poor, fire-prone, Mediterranean-type regions with a prolonged hot, dry season and exposed to intensive pressure from pollinators, herbivores and granivores are characterised by high species richness and endemism (Cowling *et al.*, 1996) and should provide suitable scenarios to examine these issues. *Hakea* is a shrub genus of over 150 species, spread throughout Australia but best represented in mediterranean southwestern Australia, and renowned for its great variation in leaf and fruit morphology, pollinators, climate and fire tolerances and susceptibility to herbivores and granivores (Groom and Lamont, 1996a, 1997, 2015; Lamont *et al.*, 2015, 2016; Hanley *et al.*, 2009, Rafferty *et al.*, 2010).

*Hakea* is highly sclerophyllous with needle-leaved and broad-leaved species. Needle leaves are twice as thick as broad leaves implying that they have had greater exposure to drought and heat (Lamont *et al.*, 2015). Previous molecular analysis indicates that *Hakea* originated in the early Miocene directly from non-fireprone, rainforest ancestors (Sauquet *et al*, 2009; Lamont and He, 2012). *Hakea* most probably originated in the sclerophyll shrublands of southwestern Australia where it continued to diversify strongly until the present (Lamont *et al.*, 2016). From the mid-Miocene, it gradually speciated and migrated onto recently-exposed, rocky substrates, sclerophyll forests and woodlands, and across the drier centre of Australia to the moister margins. Thus, we hypothesise that the ancestral *Hakea* leaves were broad, reflecting their mesic heritage, that they were retained (or re-evolved) in temperate environments, but that needle leaves arose in the mid-Miocene and proliferated strongly through the late-Miocene to present. Many hakeas have spiny leaves with a sharp apex and/or acute, marginal teeth (Barker *et al.*, 1999). Spiny hakea leaves are more effective at deterring herbivory by kangaroos than broad leaves (Hanley *et al.*, 2007), and are moderately effective at deterring black cockatoos from reaching the woody fruits of hakeas that contain highly nutritious seeds (Groom and Lamont, 2015). Macropods appeared from 17 Ma (Prideaux and Warburton, 2010), soon after the evolution of *Hakea*. The median stem of black cockatoos (Cacatuidae, Calyptorhynchinae) is positioned at 21.5–15 Ma (White *et al.*, 2011). Needle leaves lend themselves to termination by a sharp apex, so once they appeared selection pressure from vertebrate herbivores/granivores would have promoted the evolution and stabilization of sharp-tipped leaves among vulnerable lineages.

Pollinator-driven speciation has been invoked to explain plant richness in some biodiversity hotspots, since pollinator shifts usually provide effective barriers to gene flow, thereby contributing to the origin of new plant lineages (van der Niet *et al.*, 2014). Hanley *et al.* (2009) concluded that insect pollination was ancestral in *Hakea* followed by repeated bouts of bird pollination. From their molecular phylogeny of 51 *Hakea* species, mainly from eastern Australia, Mast et al. (2012) concluded the reverse. Either interpretation is possible since it is now known that honeyeaters (Meliphagidae) originated 23.5 Ma, though they only radiated strongly from 15 to 5 Ma (Joseph *et al.*, 2014). We hoped to resolve this disagreement by adding more West Australian species to our analysis. We expected bird pollination to stabilize quickly as Toon *et al.* (2014) showed for bird-pollinated legumes that this was an irreversible process.

Hakeas produce woody follicles that vary greatly in size (20–40450 mg) (Groom and Lamont, 1997). Their seeds are highly nutritious (Groom and Lamont, 2010). By far the most destructive granivore of hakeas in southwestern Australia is Carnaby’s black cockatoo (*Calyptorhynchus latirostris*, Stock *et al.* 2013). Two ways of dealing with granivores have been observed: small, camouflaged fruits that are difficult to detect visually and large, exposed fruits that resist attack mechanically (Groom and Lamont, 1997). Thus, we hypothesize that fruit size has taken two directions in *Hakea*: fruits have become either smaller (and protected or cryptic within spiny foliage, Hanley *et al.*, 2009) or larger and woodier and that these transitions are strongly unidirectional.

Recurrent fire is one of the key factors to high species richness in fire-prone ecosystems (Cowling *et al.*, 1996; Simon *et al.*, 2009). Whole-plant responses to fire can be placed into two regeneration syndromes: resprouters that survive fire and recover via dormant buds or meristems protected beneath the bark of trunks or underground organs (Clarke *et al.*, 2013), and nonsprouters that are killed by fire and population regeneration relies solely on seedlings. Both trait-types are well represented among hakeas (Groom and Lamont, 1996b). Nonsprouters live for a shorter time and have a higher fecundity than resprouters (Lamont & Wiens, 2003; Pausas & Verdu, 2005), and are therefore hypothesized to have higher speciation rates (Wells, 1969). Resprouters have an adaptive advantage over nonsprouters when a) fire is either frequent, rare or highly stochastic, b) conditions do not favor growth or seed production e.g. infertile soils or intense competition (fertile soils, high rainfall), and/or c) conditions do not favor seedling recruitment or adult survival (Ojeda, 1998, Lamont *et al.*, 2011). As the climate became more seasonal through the Miocene, the accumulation of dry matter would have been promoted and fires would have become less stochastic and more likely to occur within the lifespan of the shorter-lived nonsprouters, essential for their promotion (Enright *et al.*, 1998a) but not necessarily at the expense of resprouters (Enright *et al.*, 1998b). Increasing occurrence of arid periods (glacials) would have created mosaics of deep sands and exposed laterites and granites (Glassford and Semeniuk 1995) favoring nonsprouters and resprouters respectively (Lamont and Markey, 1995). Thus, existing knowledge does not enable us to predict the ancestral regeneration strategy of *Hakea*. Whatever trait was ancestral we can speculate that some of its descendants must soon have transitioned to the other trait, and transition and stabilization rates in both would have been similar throughout the Neogene-Quaternary.

Serotiny (on-plant seed storage) is adaptive under conditions that restrict annual seed production (poor soils, low rainfall) in fire-prone environments (for cueing seed release) with a reliable wet season (for effective seedling recruitment) (Lamont *et al.*, 1991, Cowling *et al.*, 2005). While plant death from drought is sufficient to induce seed release among hakeas, postfire conditions are still required for optimal seedling recruitment (Causley *et al.*, 2016). Thus, we postulate that strong serotiny is the ancestral condition in *Hakea* and that stabilization will be the main evolutionary process for this trait in its subsequent diversification in response to intensifying fire and seasonality. Loss of serotiny will be a later development corresponding to the gradual appearance of fire-free habitats. Hakeas that migrate to summer-rainfall grasslands (savannas) that developed in the late Miocene can also be expected to become nonserotinous.

In this study, using a time-based phylogeny for *Hakea* we assembled (El-ahmir *et al.*, 2015), we reconstructed the ancestral traits for six attributes (with 15 trait states) and determined their evolutionary trajectories in response to the advent or increasing presence of fire, seasonality, aridity, nectar-feeding birds and vertebrate/herbivores/granivores. We attempted to identify traits of the putative ancestor and the relative contribution of transition and stabilization processes to the frequency of alternative traits over geological time to account for trait representation among the extant species. Our objective was to evaluate the impact of changing environmental conditions on trait evolution and their contribution to diversification in *Hakea* to provide insights on the factors and processes explaining high species richness in this prominent Australian genus.

## MATERIALS AND METHODS

### Phylogenetic reconstruction

We built a time-based *Hakea* phylogeny (El-Ahmir *et al.*, 2015). Briefly, we included 82 *Hakea* species, each with eight gene sequences extracted from NCBI (Mast *et al.*, 2012), combined with new sequences that we generated. The outgroup included *Grevillea juncifolia, Finschia chloroxantha* and *Buckinghamia celsissima* and their DNA sequences were obtained from NCBI. We set the calibration point for the origin of the subfamily Grevilleoideae (to which *Hakea* belongs) at 70.6 Ma based on the fossil *Lewalanipollis rectomarginis* used by Sauquet *et al.* (2009). We used BEAST v2.1.0 to estimate phylogeny and divergence times under a strict clock model (Drummond *et al.*, 2006), and further details on the methods are provided in El-Ahmir *et al.* (2015).

### Trait data

We collated leaf shape and spinescence from Barker *et al.* (1999), Hanley *et al.* (2009), personal field observations and database of the State Herbarium of Western Australia (http://www.flora.sa.gov.au). Needle leaves were recognized as rounded in cross-section with a length:width ratio of >20:1. Heteroblastic species, with seedling leaves initially broad becoming needle by the end of the first growing season or seasonally broad to needle (Groom *et al.*, 1994a), were also identified. Blunt leaves had a mucro or marginal teeth with length:width ratio <1:1 while sharp leaves were >2:1.

For pollinator types, Hanley *et al.* (2009) showed that stigma–nectary distance (SND) in *Hakea* is a reliable predictor of pollinator class (also adopted by Mast *et al.* 2012). All known or putative insect-pollinated species have a SND <13 mm and all known or putative bird-pollinated species have a SND >13 mm. This is supported by the shortest bill length of the principal bird pollinators in Australia (family Meliphagidae) is 12 mm (Paton and Ford, 1977) while no known insect pollinator in Western Australia can touch the nectary and pollen presenter simultaneously if the SND >12 mm. We therefore assigned species with SND <13 mm to the insect-pollination class and >13 mm to the bird-pollination class. We took SND from Hanley *et al.* (2009) and Mast *et al.* (2012). Approximate SND for the remaining species were obtained from pistil lengths in Barker *et al.* (1999).

Fruit size, as dry fruit weight, was obtained from Groom and Lamont (1997). If not available there, the three mean fruit dimensions were obtained from Barker *et al.* (1999), converted to volume and multiplied by mean fruit density in Groom and Lamont (1997). They were divided into four size classes: <1, 1–5, >5 g, such that the 1–5 g class accounted for about half of species. Postfire response/regeneration strategy was collated from Groom and Lamont (1996b), Barker *et al.* (1999) and Young (2006). Each species was assigned as either a nonsprouter or resprouter (with two species recognized to have both fire response forms in different populations). Level of serotiny was obtained from Groom and Lamont (1997) and images on the web (especially http://www.flora.sa.gov.au).

### Trait reconstruction through the phylogeny

We used MultiState in BayesTraits (Pagel and Meade 2006) to determine the most likely ancestral traits for the *Hakea* phylogeny. First, we tested which of the possible models (simple or complex, associated with uniform rates of 0~30) should be used via the log Bayes factor (log BF) recommended by Pagel and Meade (2006). We excluded morphological data for the outgroup in order to avoid potential biases in trait assignment because they do not adequately represent the associated clades. We applied the best-fit model parameters to our MC tree in a Bayesian framework using MCMC sampling to search for optimal parameter estimates. The MCMC parameter searches consisted of 1,000,000 iterations with 25,000 discarded as burn-in. We used maximum likelihood parameter estimates as starting values in the MCMC analyses. We also used the continuous random walk (Model A) associated with the MCMC method to determine whether pairwise traits evolved in a correlated manner, and BayesFactor was used to determine the significance of correlation between any two traits (Pagel and Meade, 2006). Trait reconstruction of fruit size was carried in Mesquite using a parsimonious procedure (Madison *et al.*, 2007).

### Speciation and trait proliferation rates

Net species diversification rate was calculated as (*N*_*i* + *t*_ − *N*_*i*_)/(*N*_*i*_.*t*), where *N* is the number of lineages at the start, *i*, and end, *i* + *t*, of the time interval, *t* (He *et al.* 2011) for the three geological periods/epochs in which *Hakea* has been recorded as well as overall. The geological boundaries were set according to the International Commission on Stratigraphy (www.stratigraphy.org), while the start time in the Miocene was set at the time that *Hakea* first appeared. Following trait assignment to each node of the phylogeny, trait stabilization and transition rates (see Introduction) were determined for the three periods/epochs and overall by counting their number in each time interval. Where the ancestor was ambiguous this event was omitted from the counts as the process was unclear. They were then converted to the fraction that each process contributed to total proliferation within the trait and between all traits of that character. The number of reversals was also noted: i.e. a trait reverting to its immediate preceding trait. Individual speciation promotional rates (SPR) for the three geological periods were determined from the percentage of events retaining the ancestral trait, Y, where SPR = 100/Y (see Introduction). Generally, 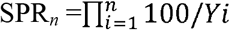 where Ys for the *n* attributes assessed are multiplied to give their total promotional effect on species diversification. SPRn was converted to its fractional contribution to species diversification for the *n* attributes assessed: (SPR_*n*_ − 1)/ SPR_*n*_.

## RESULTS

### Hakea time-based phylogeny

The Bayes MCMC analysis indicated that the *Hakea* stem arose 18.0 Ma [with the 95% highest density probability (HDP) at 15.8-20.2 Ma] and split into two clades (defined as clades A and B in Mast *et al.* 2012) 14.1 Ma (95% HPD, 12.5-15.8 Ma). The phylogeny was strongly supported by the branch posterior probability where 48 out of 81 branches were ≥ 0.70. The overall topology of *Hakea* phylogeny was consistent with that in Mast *et al.* (2012). Net species diversification rate in the Miocene greatly exceeded that in the Pliocene (9.6×) and Quaternary (13.5×) and the overall rate was dictated by the Miocene rate as it was the longest period (Table S1).

### Evolutionary trajectories for two leaf attributes

Trait reconstruction showed that the most recent ancestor (MRA) had broad leaves (*P* = 0.61) that were blunt-tipped with smooth margins (*P* = 0.88) (Fig. 2). The phylogeny split into needle (A) (*P* = 0.78) and broad (B) (*P* = 1.00) clades by 14.1 Ma. Heteroblasty arose 6.9 Ma. Both clades remained blunt-tipped (*P* = 0.69, 1.00) but sharp tips emerged in one branch of the A clade 12.7 Ma. While the transition rate for needle/heteroblastic leaves exceeded that of broad leaves in the Miocene, proliferation of broad leaves accounted for 60% of the diversification events (Table S1A). Broad leaf proliferation continued (mainly through stabilization) at the expense of needle but not of heteroblastic leaves through the Pliocene and Quaternary. Overall, 65% of total proliferations were of broad leaves (mainly stabilization) with 33 reversals to broad leaves, 30% to needle and 5% to heteroblastic (mainly recent transitions), with the overall transition rate of non-broad leaves 2.6 times broad leaves. The evolution of non-broad leaves increased the overall speciation rate by 54%, greatest in the Miocene. Spiny leaves proliferated at a similar rate as non-spiny leaves in the Miocene but the rate for spiny leaves declined slightly through the Pliocene and Quaternary due to reducing stabilization but increased slightly among non-spiny leaves due to increasing stabilization (Table S1B). Reversals were negligible. The transition rate for spiny leaves was twice that for non-spiny leaves, with their advent and proliferation increasing the diversification rate by 73%.

**Figure 2.**
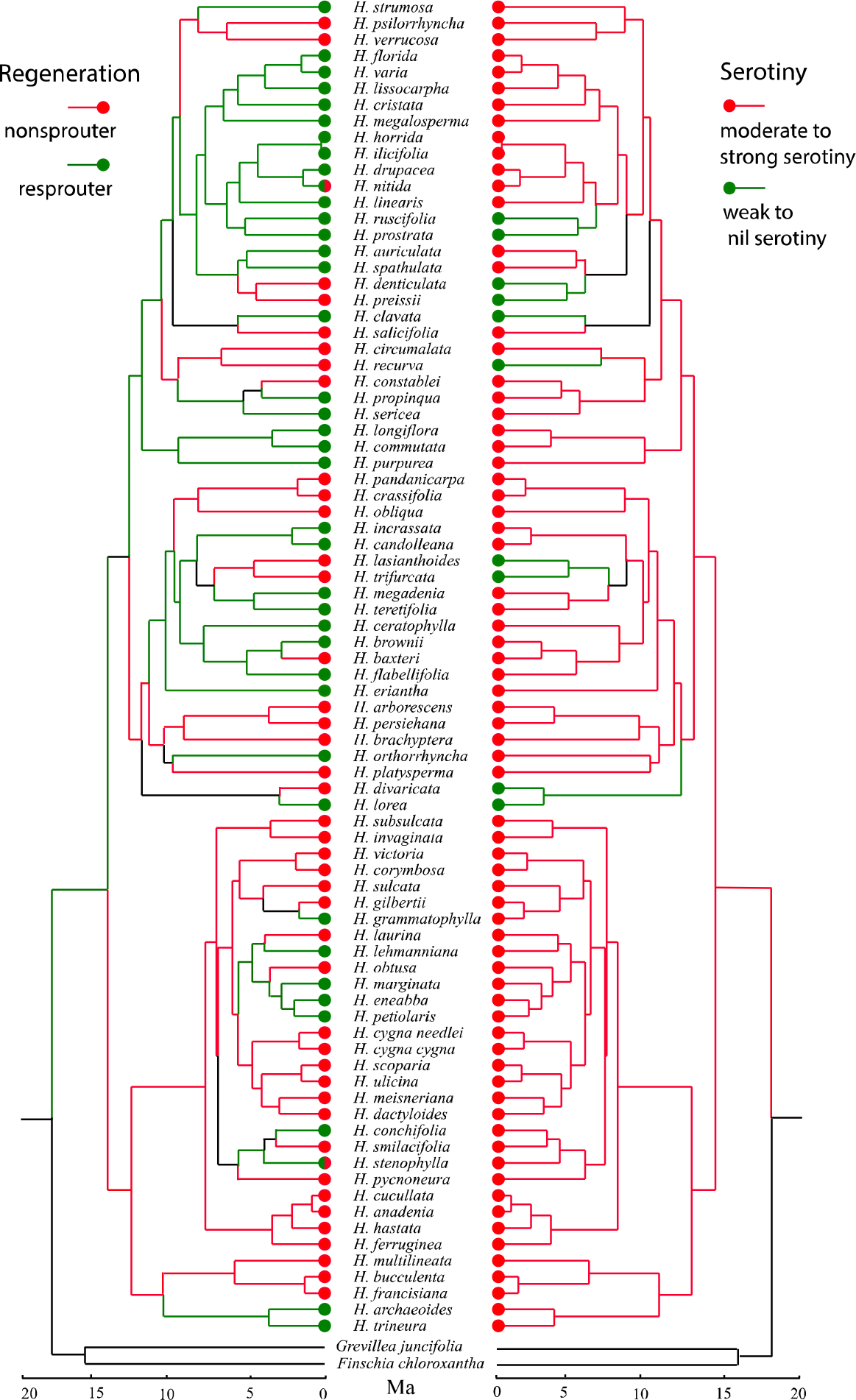
Reconstruction of leaf morphology traits through time in the genus *Hakea*. Left: leaf shape, broad, needle or heteroblastic (broad followed by needle). Right: leaf spinescence, blunt/nil or sharp apices or teeth.

### Evolutionary trajectories for two reproductive attributes

The MRA of *Hakea* showed a high posterior probability (*P* = 0.75) of being insect-pollinated. The basal split of the genus was accompanied by a shift to bird pollination 14.1 Ma in clade B (*P* = 0.82) but retention of insect pollination in clade A (*P* = 0.98) (Fig. 3). A reversal occurred in clade B 12.6 Ma while pollination transitioned to birds 12.1 Ma in clade A that remained predominantly insect-pollinated. Overall, 78 reversals occurred (Table S1C). The switch to bird pollination was restricted to the Miocene with transitions accounting for 32% of bird proliferation events, and increasing stabilization through the Pliocene/Quaternary.
Bird to insect transitions occurred in the Pliocene but not in the Quaternary. Overall transition rates for insect and bird pollination were similar, with bird pollination accounting for 30% of events and promoting speciation by 41%. The MRA had a high probability (by parsimony) of producing medium-sized fruits (1.0–5.0 g). Smaller (<1.0 g) and larger (>5.0 g) fruits first arose 12.1 Ma in clade A and smaller fruits appeared 6.5 Ma in clade B (Table S1D). In the Miocene, 19% of events involved transitions to other than medium-sized fruits but proliferation of medium-sized fruits predominated. Proliferation of small fruits (46% of events) dominated in the Pliocene, through both transitions and stabilization, and proliferation of non-medium-sized fruits contributed 150% to the stimulation of diversification events. In the Quaternary and overall, proliferation of medium and non-medium fruits contributed equally to all diversification events. Only medium fruits were sometimes the outcomes of reversals; all other transitions were unidirectional with medium→small accounting for 30 events, medium→medium-large/large for 11 events, and medium→medium-large→large for 8 events. Overall, 24% of all events involved transitions to non-medium fruits and their proliferation accounted for an 88% increase in the speciation rate.

**Figure 3.**
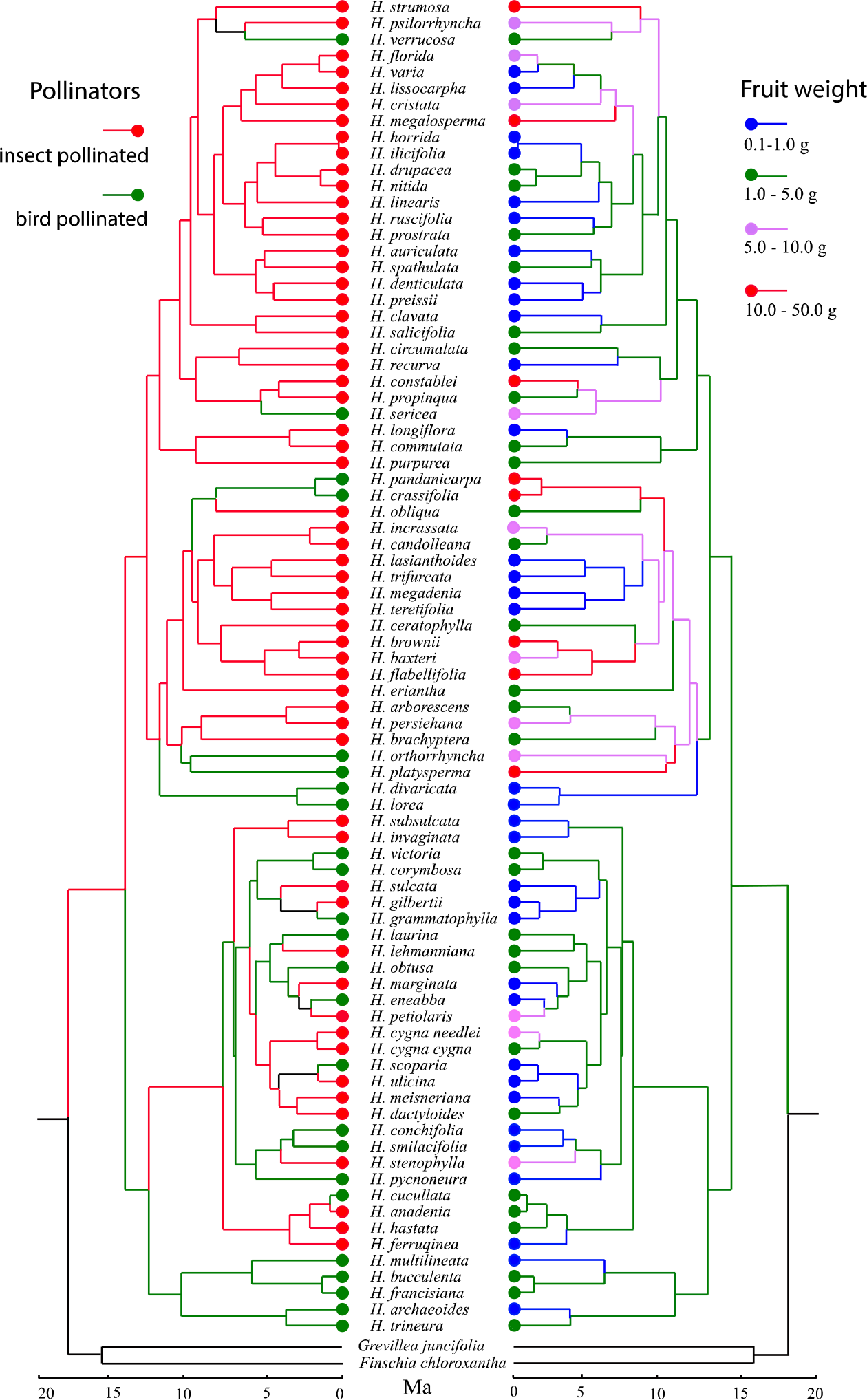
Reconstruction of reproductive biology traits through time in the genus *Hakea*. Left: insect or bird pollinated. Right: four classes of fruit size by weight.

### Evolutionary trajectories for two fire-adapted attributes

Postfire regeneration of the MRA was via resprouting though the posterior probability was not strong (*P* = 0.62). The ancestor of clade A was a resprouter (*P* = 0.73), while clade B was a nonsprouter (*P* = 0.86) (Fig. 4). By 12.7 Ma, nonsprouters also evolved in clade A. By the end of the Miocene, diversification events were spread almost uniformly between resprouters and nonsprouters (Table S1E). Transitioning to nonsprouters remained strong in the Pliocene but ceased among resprouters. Transitioning ceased in the Quaternary with nonsprouting promoting 140% more speciation through stabilization in that period. Overall, proliferation among resprouters and nonsprouters was similar with the advent of nonsprouters doubling the speciation rate due to similar high rates of stabilization, though transitions to nonsprouting approached twice that for resprouting. Reversals were common among resprouters but only 20% of reversals involved nonsprouters. Serotiny was the MRA with *P* = 1.00 and both major clades remained serotinous (*P* = 1.00). There was an isolated occurrence of weak/nil serotiny 12.1 Ma and five more subsequent origins in clade A but non-serotiny never arose in clade B. Stabilization among moderately/strongly serotinous lineages dominated trait proliferation throughout hakea's history with limited transition to weak/non-serotiny in the Miocene followed by stabilization in the Pliocene and absence of proliferation in the Quaternary.Overall, stabilization of serotiny was the main process with proliferation of non-serotiny accounting for 7% of events and it increased speciation by 8%. All transitions were unidirectional.

**Figure 4.**
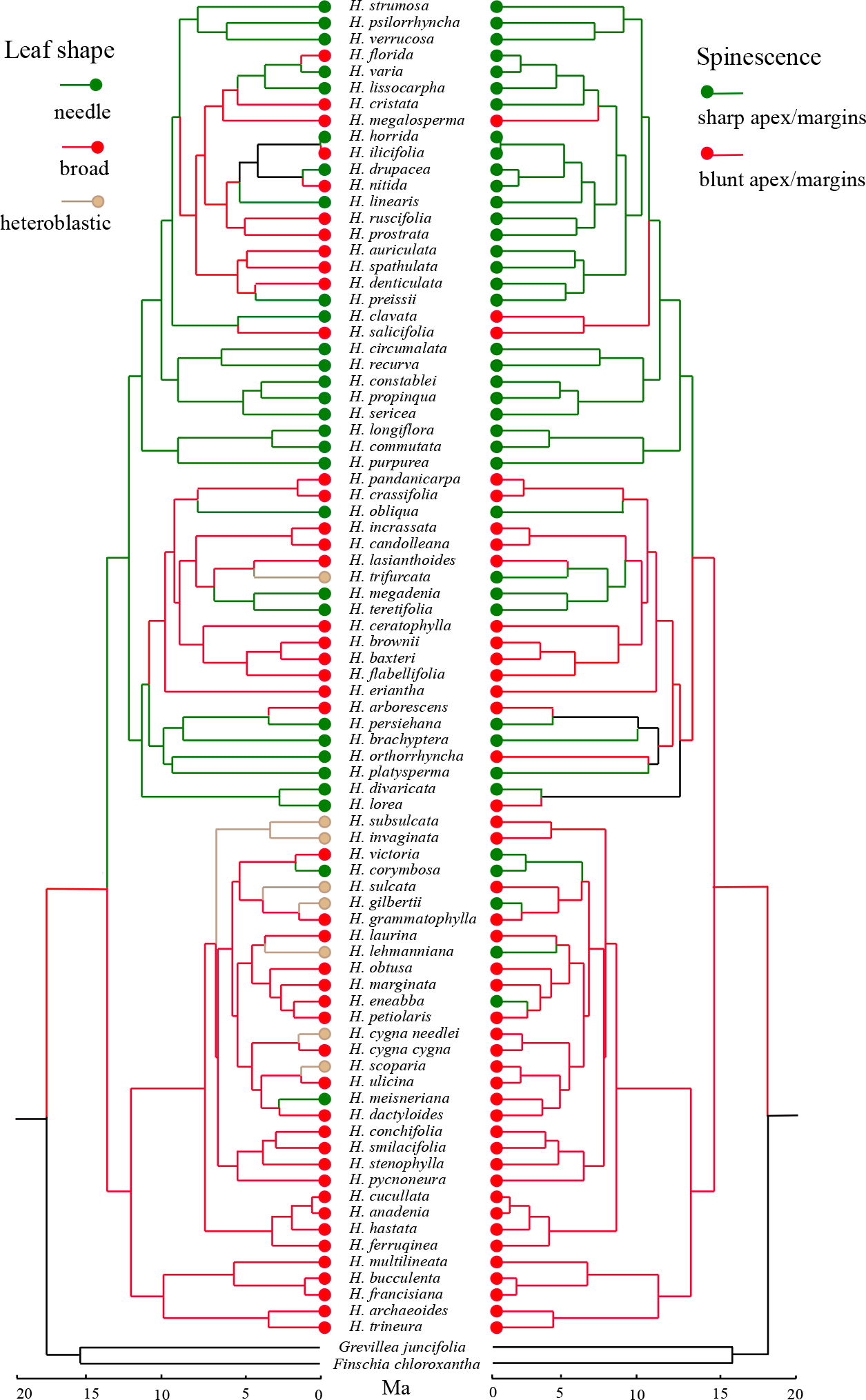
Reconstruction of fire-adapted traits through time in the genus *Hakea*. Left: nonsprouter (seedlings only) or resprouter. Right: strongly or weakly/nil serotinous.

### Promotion of species diversification

The overall speciation promotional rate (SPR6) induced by the advent of novel traits was given by 1.54 × 1.73 × 1.41 × 1.85 × 2.02 × 1.08 = 15.16. Thus, 93.4% of diversification events (ignoring reversals) can be attributed to the presence of at least one non-ancestral trait. Three species possessed the six ancestral traits (*H candolleana, H. ceratophylla, H. eriantha*), all in the same subclade of 14 species, but they included two reversals (nonsprouter→resprouter, medium-large→medium fruits). Thus, 96.3% of extant species lack at least one ancestral trait. One species (*H divaricata*) had five of six traits in the advanced condition.

### Correlated evolution between traits

Correlation analysis using the BayesFactor (BF) showed no relationship between any pairs of attributes (BF < 1.0) except leaf shape and spinescence, with needle leaves more likely to be spiny (BF = 5.3).

## DISCUSSION

Trait reconstruction of the ancestral *Hakea* phenotype shows it to have been broad-leaved, non-spinescent and insect-pollinated, with medium-sized, serotinous fruits and resprouting after fire. Resprouting and serotiny confirm that the associated vegetation was fireprone and experienced a reliable postfire wet season by 18 Ma (Lamont *et al.*, 2013). It is clear that *Hakea* changed radically at the level of fire-related adaptations, including woodiness of their fruits, when migrating from the non-fireprone environment of its ancestors (nonsprouting and nonserotinous), whereas leaf form (broad, non-spinescent) and reproductive biology [insect-pollinated, medium-sized (1-5 g) fruits] were initially conserved. Nevertheless, within 4 My, two (sub)clades had evolved with quite different syndromes of traits: one (A) that retained resprouting but possessed needle leaves many of which developed sharp apices, was bird-pollinated and where the largest woody fruits (>10 g) were produced, and the other (B) that became nonsprouting but all other attributes were dominated by their ancestral traits. The final outcome was almost equal representation of broad and needle leaves and spiny and blunt leaves, significant presence of bird-pollination, almost equal representation of small (<1 g) and large (>5 g) fruits, equal representation of resprouting and nonsprouting, and limited presence (10%) of weak/non-serotiny. Only three of 82 extant species retain all six ancestral traits and even two of these traits were the outcome of reversals. At the genus level, of 15 possible pairs of correlated evolution between attributes, only needle and sharp-pointed leaves were associated through time (attributable to their morphological links).

The species diversification rate of *Hakea* was highest by far in the Miocene than in the more recent epochs. The Miocene was a period of great climatic upheavals and the speciation rates among banksias in Australia (He *et al.*, 2011) and proteas in South Africa (Lamont *et al.*, 2013) (both genera also in Proteaceae) were also an order of magnitude higher then. The same pattern applies to proliferation of traits, with all alternative traits of the six examined highest in the Miocene (obtained by multiplying the percentage contribution to species diversification of each trait by the diversification rate on a time basis).

### Transition versus stabilization processes

While trait initiation (transition) is a vital step in speciation its incorporation into the clade (stabilization) is just as important. That proliferation of a trait through the phylogeny is rarely a function of the transition rate is strongly supported here. Taking leaf shape as an example, the transition from broad to needle leaves overall occurred at >2.6 times the rate as the reverse transition, yet stabilization of broad leaves occurred at 2.3 times the rate as needle leaves. The net result was the proliferation of broad leaves at 1.85 times the rate of needle leaves because the ratio of stabilization to transition events among broad leaves was five times the rate for needle leaves. In theory, only one initiation step is required for incorporation of a new trait into the clade provided it stabilizes quickly and is not subject to reversals. Thus, the ratio of stabilisation to transition events is a function of the strength of directional selection. The invasion of the savanna grasslands by *Protea* is a rare example of unidirectional selection associated with a single transition followed by almost universal stabilization (Lamont *et al.*, 2013). In practice, the same trait arises numerous times through the phylogeny while reversals depend on the trait. For *Hakea* leaf shape, 77% of the 43-recorded reversals were for the recovery of broad from needle leaves rendering transitions to needle less effective and reflecting unstable selective forces. The relative contribution of transition and stabilization events to proliferation depends on both the trait and the time period under consideration.

### Evolutionary trajectories for leaves

By the time *Hakea* separated from its non-fireprone ancestors 18 Ma, Australia (as much of the world) was experiencing declining levels of rainfall, temperatures and metabolically active atmospheric gases, and increasing seasonality. In addition, the opening up of the vegetation would have exposed them to high light intensity and diurnal temperatures (Jordan *et al.*, 2005) compared with closed forests. If needle leaves increase fitness to such constraints, and currently they account for 45% of species so this genus has a strong propensity to produce them, they should have evolved early in its history and proliferated through stabilization. Indeed, within 4 My, a needle-leaved clade (A) had arisen with strong stabilization leading to 56% of its extant species being needle-leaved. Evolution of needle leaves was greatly delayed in clade B and was mainly expressed through the appearance of heteroblastic species over the last 5 My (all from broad-leaved ancestors). The latter appeared so recently that there have been no opportunities for reversals unlike needle leaves where reversals to broad have been frequent. These reversals confirm the lability of leaf form among isobilateral leaves as demonstrated ontogenetically by *H. trifurcata* whose juvenile leaves are needle, a few becoming broad at the start of the growing season in adult plants and needle again as the dry summer approaches (Groom *et al.*, 1994a).

The dominance of broad leaves and reversals to them require some explanation. Clearly, leaf form is not the only way of dealing with drought, such as deep root systems (Groom and Lamont, 2015), while broad leaves among hakeas are still highly sclerophyllous (Lamont *et al.*, 2015) with thick cuticles, sunken stomata and a tannin-filled hypodermis (Jordan *et al.*, 2005). They are often narrow or strap-shaped rather than truly broad, such as *H. grammatophylla* in the 'deadheart' of Australia, and all are vertically oriented. In addition, broader-leaved species have retreated to the moister parts of the landscape or subregions (Groom and Lamont, 1996a). In fact, the frequent reversals in both directions are consistent with climatic oscillations that became characteristic of the Pliocene/Quaternary and their evolutionary tracking.

While broad leaves may be spinescent, such as *H. cristata*, needle leaves that are already rigid and with a sclerified apex can readily be transformed into strongly piercing structures through elongation and thinning of the mucro. This morphogenetic link explains the unique evolutionary correlation through time of needle and spinescent leaves. Thus, following a small delay, one branch of the A clade became spinescent at 12.7 My. The broad-leaved B clade remained essentially non-spinescent. If spinescence is effective against herbivores (macropods) and florivores/granivores (emus, cockatoos) (Hanley *et al.*, 2007) the delay in its appearance cannot be attributed to their absence as all were present by this time but they speciated gradually and their selective effects would have intensified over time. It is of interest that transitioning to spiny leaves was most marked in the Quaternary, a time when modern cockatoos evolved in SW Australia, though their ancestors were present from the early Miocene (Joseph *et al.*, 2014). Reversals were negligible indicating strong directional selection. Why more events did not yield spiny leaves is partly attributable to morphological constraints, the fact that all *Hakea* leaves are highly unpalatable and not grass-like (Rafferty *et al.*, 2010), and ability of vertebrates to learn to overcome physical deterrents (Hanley *et al.*, 2007).

### Evolutionary trajectories for flowers and fruits

Bird-pollinated flowers evolved from insect-pollinated flowers with the split of the genus 14.1 Ma. This resolves the disagreements over which was the ancestral condition (Hanley *et al.*, 2009; Mast *et al.*, 2012) caused by misidentifying the basal lineages or not including sufficient (representative) insect-pollinated lineages from SW Australia where the clade most probably arose (Lamont *et al.*, 2016). Honeyeaters (Meliphagidae) were already present in Australia at the time *Hakea* originated, but these birds only diversified strongly in the mid to late-Miocene, especially among such major pollinators as *Phylidonyris, Anthochaera, Lichmera* and *Lichenostomus* (Joseph *et al.*, 2014). In fact, apart from *H. cucullata* in the Quaternary, the only time flowers increased their size to accommodate bird pollinators was in the Miocene. For the Pliocene/Quaternary it was stabilization processes only. The greater levels of stabilization among insect-pollinated lineages throughout their history explains their current greater abundance and suggests that they have been a greater selective force, possibility associated with their greater reliability rather than morphological diversity that would have favored greater transition rates.

Of note are the transitions from bird to insect pollination in the Miocene/Pliocene and the large number of reversals (78), 60% of which were insect**→**bird**→**insect. This is significant on two counts: trait reversibility and fluctuating selection. Bird-pollination is regarded as an evolutionary 'dead-end’ because specialization of floral structures for birds may be irreversible (Toon *et al.*, 2014) so shifts from bird to insect pollination are rare (van der Niet & Johnson 2012). This hypothesis derives from hummingbird pollination systems that require specialised floral structures (Tripp and Manos, 2008). However, specialised floral structures are not essential for honeyeaters because they are generalist pollinators and not obligate nectarivores. This lack of specialization in the honeyeater pollination system implies minimal floral structure specialization – simple elongation of the pistil is sufficient (Hanley *et al.*, 2009). As a result, reversal shifts from bird to insect are possible in situations when bird pollinators become scarce. The evolutionary tracking of the great climatic fluctuations, with their profound effects on the abundance of both birds and insects, that occurred from the mid-Miocene can explain the remarkable number of reversals in both directions. Such great flexibility in pollinator shifts may provide insights in explaining the mechanisms that promoted explosive speciation in *Hakea* but even more so in its younger sister, *Grevillea* with 150 of 362 species being bird-pollinated (Ford *et al.*, 1979).

Morphological variation in fruit structure among hakeas is exceptional and transition rates away from the ancestral medium-sized ancestor equalled their stabilization rates through each of the three epochs. Two groups of granivores, insects and parrots, were already present when *Hakea* emerged. The transition to small (camouflaged) fruits, which appear better protected against insects than cockatoos (B Lamont, unpublished), was strongest in the Miocene and continued throughout the period, by far the highest among all traits we assessed. Large woody fruits, particularly effective against cockatoos (Groom and Lamont, 2015), followed a similar path. There were no reversals among small and large fruits indicating strong unidirectional selection for these extremes.

### Evolutionary trajectories for fire-related traits

The twin ancestral traits of resprouting (adaptive in the presence of severe, periodic disturbances where recruitment opportunities are limited) and serotiny (ensuring seeds are released when those limited recruitment opportunities are optimal) demonstrate that *Hakea* arose in a fireprone environment. We can also surmise that fires were of overall moderate frequency (>5-45 y intervals) but highly stochastic. If fires were at the high frequencies associated with savanna grasslands (<5 y intervals) the plants would have stayed nonserotinous (Lamont *et al.* 2013). If fire intervals exceeded their longevity then, upon death in the absence of fire, serotinous seeds would have been released onto a hostile seedbed and rarely yielded recruits (Causley *et al.*, 2016). However, within 4 My, a nonsprouting clade (B) had arisen. The outcome was strong transitioning to both fire-response types in the Miocene and all epochs were dominated by stabilizations against a background of almost universal proliferation by serotinous descendents. Of note are both the continuing transitions to nonsprouting in the Pliocene and its steady increase in stabilization rate throughout *Hakea*'s history. It is likely that the trend of increasing aridity and seasonality and declining atmospheric oxygen and carbon dioxide (He *et al.*, 2012) led to less frequent, but more reliable, fires and promotion of nonsprouting (Lamont *et al.*, 2011, 2013). Today, resprouters are better represented in the more fireprone, strongly seasonal northern sandplains of SW Australia and nonsprouters in the less fireprone parts, especially on deeper soils where recruitment and adult survival are more likely (Lamont and Markey, 1995; Groom and Lamont, 1996a).

Transitions to non/weak serotiny were rare but five of the six independent origins were in the late Miocene and proliferation was restricted to the Pliocene. Explanations vary but include migrations to frequently-burnt savanna grasslands (*H divaricata* lineage) or rarely-burnt aridlands (*H. recurva*), exposure of novel firefree rock outcrops to which some species adapted (*H. clavata*), and presence in forests with reliable winter rains ensuring recruitment interfire (*H. trifurcata* lineage) (Hanley and Lamont, 2001). This pattern has limited parallels with *Protea* in South Africa where one transition to nonserotiny in grasslands was followed by increasingly extensive stabilization there that failed to occur in Australian grasslands. However, in both super-regions stabilization of the ancestral condition, serotiny, was by far the dominant process and reversals were negligible. By contrast, the well known lability in whole-plant fire responses (He *et al.*, 2011) was expressed in both *Protea* and *Hakea*, although 80% of reversals were to resprouting in *Hakea* in the Miocene, perhaps reflecting periods of increased or more stochastic fire frequencies as the clade moved to other parts of Australia. The historic levels of fire frequency coupled with the severity of seasonal droughts serve well to interpret the relative abundances and distributions of resprouters and nonsprouters among hakeas, ericas and proteas (Ojeda, 1998; Lamont *et al.*, 2013).

### Promotion of speciation by non-ancestral traits

There are two ways of considering the role of traits in speciation. One is to compare how they have *contributed to* all diversification events and the other is to estimate to what extent non-ancestral traits have *promoted* these diversification events. The latter is hypothesis-driven and based on the premise that without trait innovation and subsequent stabilization these diversification events would not have occurred. Thus, it relies on being able to identify both the ancestral state and its pathway through the phylogeny in order to ascertain trait reversals. This means that any extinction events cannot be incorporated into the analysis, but we argued earlier (see Introduction) that most new traits are adjunct to (as they are spatially displaced), rather than replace, ancestral traits anyway. Nevertheless, entire but unknown lineages characterized by certain historically maladapted traits may be missing (or only represented by long branches in the chronogram) if there have been radical environmental shifts that no amount of adjustment, in the absence of fossil evidence, can correct for. However, there is little evidence of extinctions as a significant evolutionary process in the SW Australian flora over the last 10 My (Hopper, 2009).

For the six *Hakea* attributes examined, proliferation of the non-ancestral trait promoted speciation by 1.08 (weakly serotinous) to 2.02 (fire-killed) times. Overall, this increased speciation by 15.2 times, equivalent to 93.4% of diversification events. Two of the ancestral traits possessed by the three species of identical phenotype to the putative original phenotype were the result of reversals so that even these were the outcome of trait diversification when the ancestral condition would have been temporarily lost. Reversals were particularly prevalent among pollination types and fire-response types but absent altogether from the serotiny types. They represent the net effect of a) the constancy of directional selection, b) the lability of opposing traits, and c) time available for further transitions to occur. Certainly, the high level of lability among fire-response types is consistent with previous studies on *Protea* (Lamont et al. 2013), *Banksia* (He *et al.*, 2011) and Restionaceae (Litsios *et al.*, 2014).

Clearly, *Hakea*’s remarkable genetic/morphological malleability in the face of these strong selective agents has resulted in an exceptionally diverse clade and its distribution throughout Australia. We might now wonder to what extent the cumulative contributions by transitions and stabilizations to trait proliferation among critical plant attributes, in response to an array of environmental constraints introduced in the Miocene, have led to the explosive radiation of such speciose genera as *Grevillea, Acacia, Melaleuca* and *Eucalyptus*, that currently dominate Australia's sclerophyll flora, and the floras of many other parts of the world subject to similar selective forces.

## Acknowledgments

This study was supported by the Australian Research Council (DP120103389 and DP130103029).

---

**Table S1.**
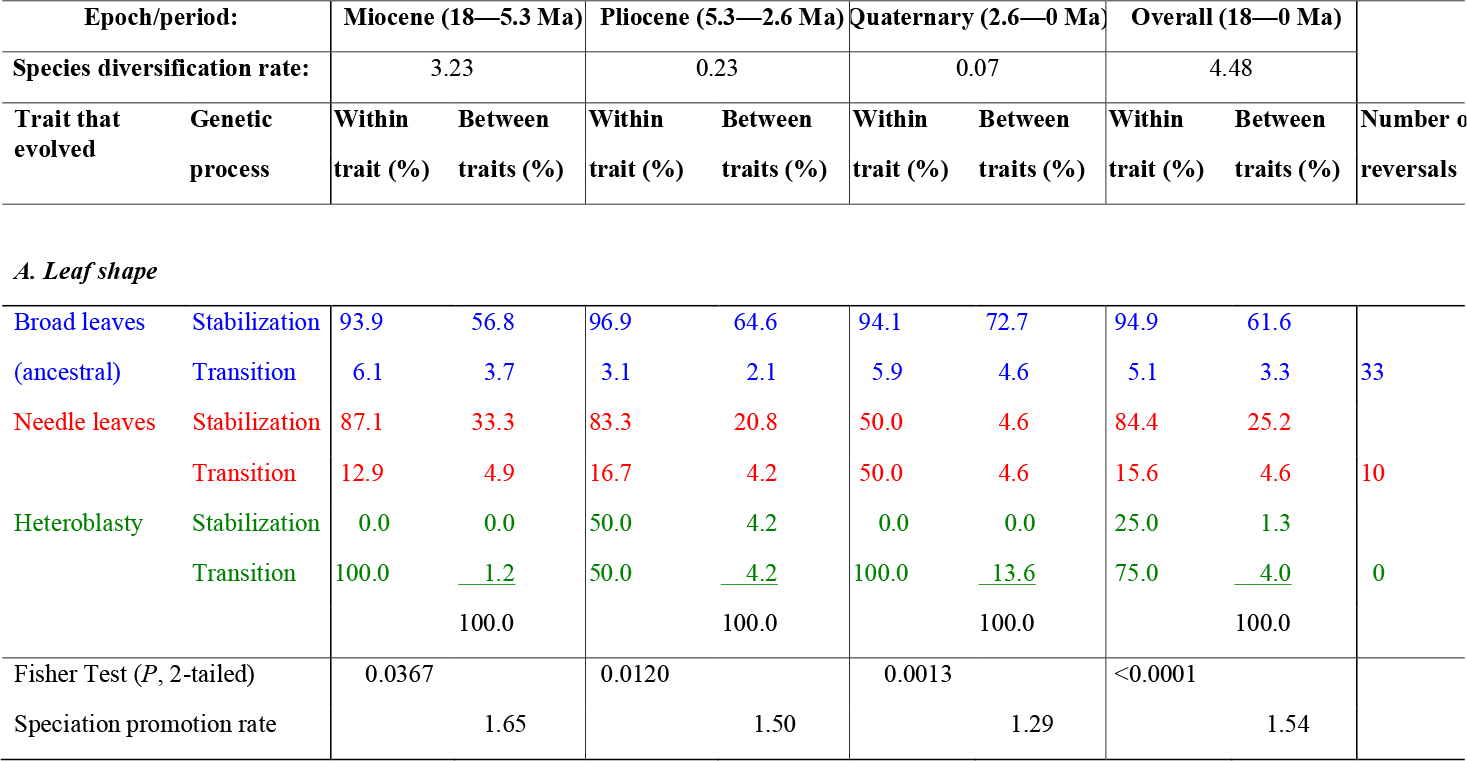
Paired trait evolution in *Hakea* apportioned among stabilization (trait retained during diversification event) and transition (trait attained during diversification event) processes in each Epoch based on the molecular chronogram reported here. All node-to-node steps in the phylogeny were treated as diversification events. Where the ancestor was ambiguous this event was omitted from the counts as the process was unclear. Reversals refer to transitions back to the previous trait.

